# A new Critically Endangered cloud forest tree *Microcos* (Grewiaceae-Malvaceae) from the Rumpi Hills of S.W. Region Cameroon

**DOI:** 10.1101/2022.08.03.502699

**Authors:** Martin Cheek, Sara Edwards, Jean Michel Onana

## Abstract

We describe *Microcos rumpi* (Grewiaceae-Malvaceae) as a new species to science from the Rumpi Hills of SW Region Cameroon, a proposed Tropical Important Plant Area. Confined on current evidence to submontane forest, the species is threatened by expanding habitat clearance for farms and is assessed as Critically Endangered. A massive tree, attaining 35 *–* 40 m height, and 80 cm trunk diameter at 1.3 m above ground, its biomass is calculated as in the range of 7 *–* 8 metric tonnes. It is the third tree species of the genus recorded from Cameroon and only the fourth recorded west of D.R. Congo. A key to these four species is presented.

The concept of *Microcos* in Africa in relation to *Grewia* is discussed, and three new combinations are made, transferring three species names from *Grewia to Microcos*: *Microcos louisii* (Wilczek) Cheek, *M. evrardii*(Wilczek) Cheek and *M*.*schmitzii* (Wilczek) Cheek.

## Introduction

In connection with the Cameroon Tropical Important Plant Areas programme (Darbyshire *et al*. 2017; Cheek, continuously updated), in May 2022, while clearing a backlog of specimens which had resulted from a survey of the Rumpi Hills and Nta Ali in S.W. Region Cameroon (Thomas 1995), two specimens, *Thomas* 10426 and 10442 (both K!) that had been field-identified as *Grewia coriacea* Mast. (more correctly, see below, *Microcos coriacea* (Mast.) Burret), were found to be discordant with that species. Further checks against reference specimens in the Kew herbarium showed that not only did they differ from *Microcos coriacea* but from all other *Microcos* species in tropical Africa (see Results below). Accordingly, these Rumpi Hills specimens are here formally described and named as *Microcos rumpi* Cheek.

*Microcos* Burm. ex L. (1753) is a palaeotropical genus of about 78 species (Govaerts et al., continuously updated) based on *M. paniculata* Burm. ex L. (1753) from Sri Lanka. Linnaeus (1767) later synonymised *Microcos* under *Grewia* L. However, the genus was resurrected by Burret (1926). Burret’s authoritative revision (1926) of former Tiliaceae sens. lat. Presaged its break-up into todays’s Brownloideae/Brownlowiaceae, Tiliaceae sensu stricto/Tilioideae and Grewioideae/Sparrmanniaceae (Grewiaceae) including *Microcos* (Bayer *et al*. 1999, Bayer & Kubitzki 2003, Cheek in Heywood *et al*. 2007). Burret’s was the last global treatment of *Microcos* (Burret 1926). He recognised 53 species, of which 19 were recorded from Africa and 34 in Asia. Of the 99 names in *Microcos* listed in IPNI (continuously updated), Govaerts *et al*. continuously updated) accept 78 names. The majority of these 78 are in S.E. Asia, but with 11 in Africa. *Microcos* is absent from the Neotropics and Madagascar.

Illogically, while *Microcos* has been maintained as a separate genus from *Grewia* in Asia (e.g. Chung 2003, 2006, Chung *et al*. 2005a, Chung & Soepadmo 2011), the two genera have often been united under *Grewia* in Africa. For example, in one of the most recent Flora accounts of *Grewia* (including *Microcos*) for Africa, Whitehouse in Whitehouse *et al*. (2001) states “ …Kirkup followed Burret in recognising *Microcos* as a distinct genus; this concept has also been followed in SE Asia. Although there are clear differences between *Microcos* and the other sections of *Grewia*, for consistency I am following the practice set by the other African floras, of not recognising….” This practice of retaining *Microcos* in *Grewia* is maintained widely today in Africa, for example by the excellent and essential African Plant Database (continuously updated). In fact, the two genera are readily separated as expressed in the key below, modified from that in Whitehouse (2001):

Trees and climbers, rarely shrubs, of evergreen forest; stigmas entire; fruit unlobed; inflorescences terminal, sometimes axillary also, many-flowered……**Microcos**

Shrubs, rarely trees, of bushland or woodland; stigmas lobed; fruit 4-lobed, rarely entire; inflorescences usually axillary or leaf-opposed, rarely terminal, usually few-flowered…………………………………………………………………..**Grewia**

According to the molecular analysis of Brunken & Muellner (2012), *Microcos* is not embedded in *Grewia*, neither are these two genera sisters, in fact they fall into distinctly separate clades. Additional characters for separating the two genera are found in the pollen, wood anatomy and in the leaf anatomy, particularly the epidermal cells (Chattaway 1934, Chung 2002, Chung *et al*. 2003, 2005b). *Microcos* was maintained in Bayer & Kubitzki (2003). However, in Africa, several species described in *Grewia* remain to be transferred formally to *Microcos* which is partly addressed in this paper (see below).

In contrast to Asia (see references above), the genus *Microcos* has been little studied in Africa, as evidenced by the fact that the first new name in African *Microcos* since 1926 was published in 2004 (*Microcos barombiensis* (K. Schum.) Cheek in Cheek *et al*. 2004: 414). In the course of matching the material described as new in this paper, it became clear that a revision of the genus for Africa is desirable to address specimen misidentifications and additional apparently undescribed species. It is hoped to address these problems in future.

## Methods

Herbarium citations follow Index Herbariorum (Thiers *et al*. continuously updated) and binomial authorities IPNI (continuously updated). Material was collected using the patrol method (e.g. Cheek & Cable 1997). Material of the suspected new species was compared morphologically with material of all other African *Microcos* (or *Grewia* sect. *Microcos* (L.)Wight & Arnott) principally at K and YA, but also using images of specimens available online, principally from BR, P and WAG. Burret’s types of *Microcos* at B were destroyed by allied bombing in 1943 so it was not possible to consult them. This disaster has necessitated that subsequent authors select neotypes of his names, e.g. Whitehouse (2001).

The conservation assessment was made using the categories and criteria of IUCN (2012). Herbarium material was examined with a Leica Wild M8 dissecting binocular microscope. This was fitted with an eyepiece graticule measuring in units of 0.025 mm at maximum magnification. The description terminology follows Beentje & Cheek (2003) and Cheek (2017). The drawing was made with the same equipment using Leica 308700 camera lucida attachment.

## Taxonomic Results

Here we formally transfer three more names from *Grewia* to *Microcos* for the reasons given in the discussion, and so that one of them can be referred to in the context of the taxonomic placement of the main subject of this paper, *Microcos rumpi* (see below). All three names are of taxa of D.R. Congo published in what was intended to be a precursory account to the Flore de Congo Belge Tiliaceae treatment, although the last preceded the first by several months.

### *Microcos louisii* (R.Wilczek) Cheek comb. nov

*Grewia louisii* R.Wilczek (1963a:20; 1963b: 460). Type: D.R. Congo, District Forestier Centrale, Yalulia, 20 km a l’Est de Yangambi, foret maracageuse de la riviere Butale, alt ? 470 m, petit arbre, fl. Fr. 30 May 1938, *Louis* 9549 (holotype BR barcode BR00000897140!; isotypes BR00000197143!, YBI barcode YBI154249425!)

### *Microcos schmitzii* (R.Wilczek) Cheek comb. nov

*Grewia louisii* R.Wilczek (1963a:20; 1963b: 459). Type: D.R. Congo, District du Haut-Katanga: Elisabethville (now Lumbumbasi), Muhulu, arbuste parfois lianeux, fl. fr., 20 April 1949, *Schmitz* 2288 (holotype barcodes BR0000008930415! isotypes BR00000893041!, K000241788!, KIP468350430, KIP225201638!, KIP555461425!, PRE0271994-0!, YBI155054003!)

### *Microcos evrardii* (R.Wilczek) Cheek comb. nov

*Grewia evrardii* R.Wilczek (1963a:24; 1963b: 463). Type: D.R. Congo, District Forestier Centrale, Mondombe-Yalusaka (Terr. Ikela), Ingende, foret maracageuse, arbuste, fl.fr. 12 avril 1959, *Evrard* 6116 (holotype barcode BR00000896343!; isotypes YBI104161000!) Here we present a key to the *Microcos* tree species west of D.R. Congo, a table of characters separating the three Cameroon species tree species, and the similar *M. louisii* of D.R. Congo, from each other. We then formally describe and name the new entity, *Microcos rumpi*.

## KEY TO THE SPECIES OF MICROCOS TREES IN AFRICA WEST OF D.R. CONGO

1. Leaves deeply toothed. Maiombe Mts of Cabinda……………..***M. gossweileri*** Leaves entire. Nigeria to Congo-Brazzaville, unknown from Cabinda……….2
2. Leaf base cuneate, leaf surfaces glabrous …………………………..***M. coriacea*** Leaf base truncate or cordate, leaf blade lower surface stellate hairy 3
3. Leaves with domatia; abaxial surface 10 *–* 20% covered with scattered stellate hairs; lateral nerves (7 *–*)8 *–* 9(*–* 10) on each side of the midrib; fruits smooth ……………………………………………………….***M. rumpi sp. nov***. Leaves without domatia; abaxial surface >80% covered with stellate hairs (hairs touching each other); lateral nerves 11 *–* 13 on each side of the midrib; fruits verrucate ………………………………………………….…….***M. magnifica***

### *Microcos rumpi* Cheek *sp. nov*

Type: Cameroon, S.W. Region, Korup Project Area, Rumpi Hills, near Madie River 4^°^ 58’N, 9^°^ 15’E, fr. 22 Feb 1995, *D*.*W*.*Thomas* 10426 (holotype K, barcode K000875935; isotypes EA, K, MA, US, YA). (Fig. 1).

**Fig. 1.**
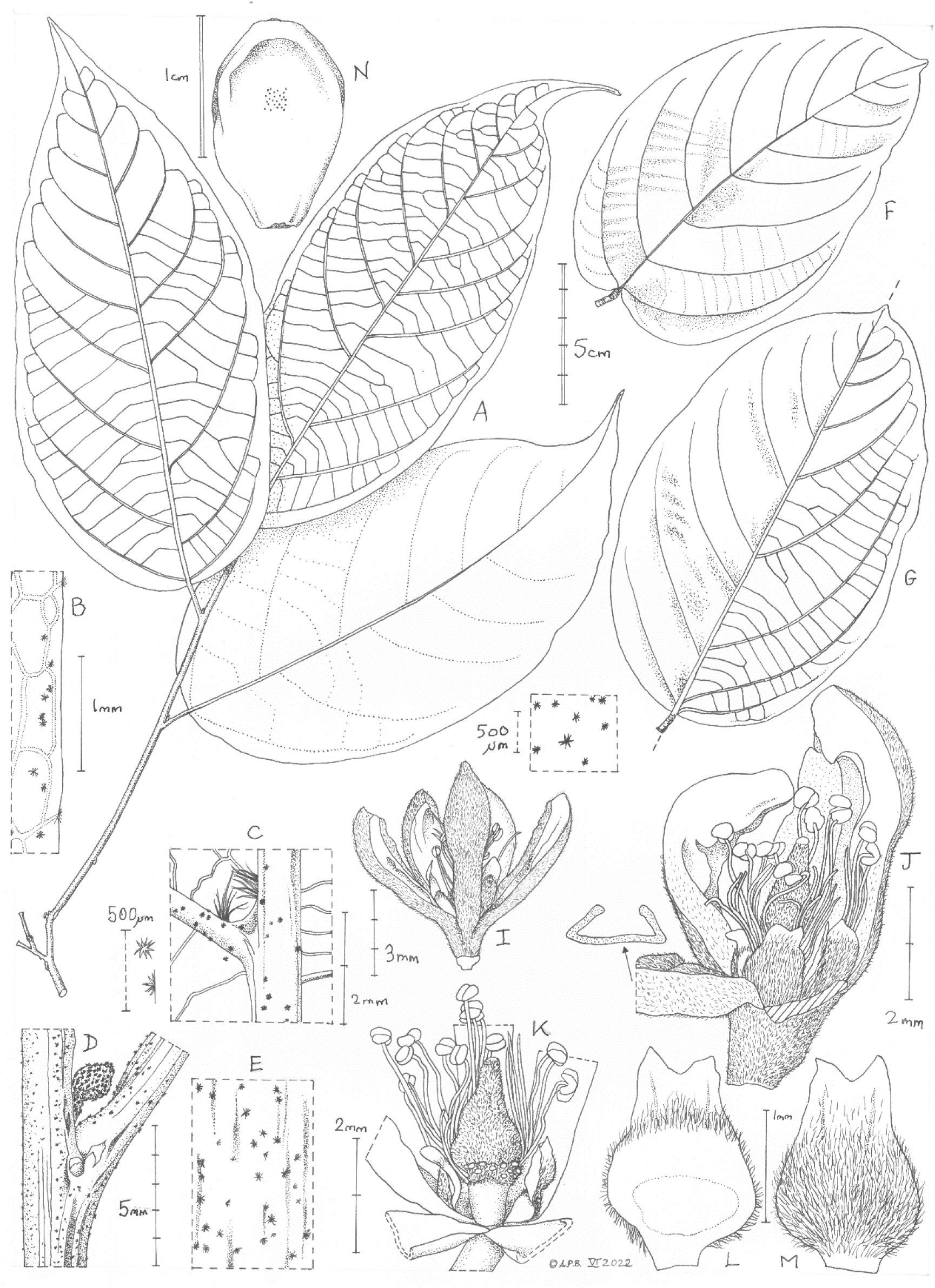
Microcos rumpi. **A** habit of leaf stem (sapling); **B** abaxial leaf margin (detail from A); **C** domatia (detail from A); **D** stem node showing stipule scar, supra-axillary bud and petiole base (detail from A); **E** indumentum on stem (detail from A); **F** mature (canopy) leaf, adaxial surface; **G** mature (canopy) leaf, showing adaxial, pleated surface on left and abaxial (scalariform tertiary nerves) on right; **H** indumentum, abaxial surface (from G); **I** flower, side view; **J** flower three sepals removed; **K** flower showing androgynophore and ovary (after removal of intervening stamens, petals and sepals); **L** petal, adaxial surface; **M** petal, abaxial surface; **N** fruit, side view. **A-E, I-M** from *Thomas* 10442(K); **F-H** from *Thomas* 10426 (K, holo.). All drawn by ANDREW BROWN.

*Grewia coriacea* sensu D.W.Thomas (1995), non Mast. (Masters 1868).

*Canopy emergent evergreen tree, 35 – 40 m tall*, and to 80 cm diam. 1.3 m above ground-level, trunk irregularly fluted at base, slash pink and fibrous looking, not peeling. Leafy stems (from saplings) terete 2 *–* 3 mm diam., internodes 1.6 *–* 3.3 cm long, epidermis purple black, smooth, 20 *–* 50% covered by a range of stellate to simple dull white to yellow hairs, stellate hairs 10 *–* 15-armed, 0.1 *–* 0.15 mm diam., simple, bifid, trifid and cruciate hairs 0.025 *–* 0.05 mm long (Fig. 1E). Previous year’s stems with surface longitudinally furrowed and with raised, longitudinally elliptic, pale brown, lenticels 0.5 × 0.2 *–* 0.35 mm. *Leaves* alternate, blades simple, entire, coriaceous, sub-bullate, drying mid brown, (ovate-) ovate-elliptic to elliptic-oblong (elliptic-obovate), (14.2 *–*)15.1 *–* 19(*–* 28.8) x (8 *–*)9.2 *–* 11(*–*11.8) cm, apex subacuminate or acumen short and broad, obtuse, 0.4 *–* 0.8(*–* 0.9) x 0.4 *–* 1.0(*–* 1.2) cm, base slightly asymmetric (broadly rounded-) truncate (-cordate), sinus c. 5 × 18 mm; abaxial surface pinnately nerved (not tri- or palmately nerved at base), lateral nerves prominent (7 *–*)8 *–* 9(*–* 10) on each side of the midrib, arising at c. 50 ° from the midrib, straight, and then arching gradually upwards, becoming parallel with the margin then uniting with the nerve above via tertiary nerves, forming looping inframarginal nerves 3 *–* 4 mm from the margin in the distal half of the leaf. Domatia conspicuous, at junction of lateral nerves and midrib with a bright white dense tuft of hairs on each, the two tuft bases separated by c. 0.5 mm, the hairs 0.5 *–* 1 mm long, the apices of the hairs of the two tufts interlocking (Fig. 1C). Tertiary nerves very prominent, scalariform, 14(*–* 17) between the basal and near basal secondary nerve pairs. Quaternary nerves barely detectable with the naked eye, reticulate, cells isodiametric, 0.75 *–* 1.25 mm diam.; indumentum of dull white mainly 6 *–* 8-armed stellate hairs 0.1 *–* 0.2 mm diam., covering 10 *–* 20(*–* 30%) of the surface; adaxial surface with nerves impressed, quaternary nerves invisible, glabrous; margin slightly revolute and thickened, entire.

*Sapling leaves* (*D*.*W. Thomas* 10442, K) as mature, canopy leaves but papyraceous, lanceolate, 20 *–* 21 × 8.9 *–* 9.7 cm, acumen narrowly triangular 2 *–* 2.3 × 0.6 *–* 0.8 cm, base rounded-truncate, lateral nerves 9 *–* 11 on each side of the midrib, quaternary nerves reticulate, conspicuous with the naked eye, finer nerves conspicuous with lens, also reticulate, cells c. 3 mm diam.,; margin slightly sinuate, with veinlets terminating in minute purple glandular teeth, teeth 3 *–* 4 mm apart, broadly obtuse, c. 0.1 × 0.15 mm, indumentum extremely sparse. *Stipules* lateral to petiole base, caducous, cicatrices white, sub-isodiametric, 0.7 *–* 1 × 1 mm. *Petiole* cylindrical (0.8 *–*)1 *–* 1.4(*–* 1.5) x 0.2 *–* 0.3 cm, indumentum as stem. *Buds* supra-axillary, inserted 0.5 *–* 1 mm above the axil, narrowly ellipsoid 1.5 × 0.75 *–* 1.2 mm, completely covered in grey stellate hairs. *Inflorescence* not seen, probably terminal, paniculate. *Flowers* (rehydrated from *Thomas* 10442) pre-anthetic buds obovoid-cylindric 7.5 *–*9 × 4 mm, pedicel 1.5 *–* 2 × 1 mm, 5 ridged, ridges rounded, minutely puberulent. *Sepals* 5, valvate, divided to the base, oblong-spatulate, 6 *–* 7 × 1.7 *–* 2 mm, the proximal half narrowed, c. 1.1 mm wide, in bud marginal ⅓ inflexed, surface minutely and densely simple puberulent (Fig. 1I&J). *Petals* 5, dark red, 2 *–* 2.2 × 1 mm, basal part ovate-elliptic 1.1 *–* 1.2 × 1 mm, distal part narrowed, oblong, 1 × 0.25 mm, sometimes retuse, outer (abaxial) surface with basal part completely covered in dense golden *–* yellow to white stellate hairs, 3 *–* 5-armed, arms erect, 0.05 *–* 0.11 mm long, distal part glabrous; inner surface with basal part divided into three portions, from base to apex: a) densely golden papillate zone c.0.5 mm long, b) dark red glabrous zone c.0.3 mm long, c) densely long-white hairy zone c.0.4 mm long, with hairs 0.05 *–* 0.07 mm long, extending along the margins (Fig. 1L&M); distal part more or less glabrous but with a few white stellate hairs at base. *Androgynophore* yellow, longitudinally 5-angular 1 × 1.1 mm, glabrous; surmounted by a subrugose platform 0.5 mm long, 2.25 *–* 2.4 mm wide, densely short simple hairy (Fig. 1K). *Stamens* c. 60, free, in c. 3 whorls, completely concealing the ovary, filaments cylindrical, dark red, glabrous, 2.1 *–* 6 mm long, the shortest outermost; stamens dithecate, medifixed, subhemispherical, 0.5 mm diam. *Ovary* ovoid-conical, 1.75 × 1.25 mm, with c. 6 shallow, longitudinal, rounded ribs, apex obtuse-rounded, densely pale yellow simple puberulent; style stout, black, 0.3 mm diam., tapering from base to apex. Stigma white, capitate-discoid, 0.1 *–* 0.12 mm diam. Surface minutely papillate. *Fruits* brown or purple-brown, glossy, obovoid 3.3 *–* 5 × 2 *–* 3.2 cm, pericarp hard, brittle, 0.25 mm thick (Fig. 1N); mesocarp with long, soft, dull pale yellow dense fibres, each c. 0.05 mm diam. Endocarp ovoid 9 *–* 12 × 6 *–* 7 mm, pale brown, the outer surface smooth, with lines of fibres attached, 2 *–* 3-valved, valves longitudinal, c. 0.5 mm thick. *Seed* narrowly ovoid 8 *–* 9 × 5 mm, pale brown.

## RECOGNITION

*Microcos rumpi* Cheek differs from *M. coriacea* (Mast.) Burret in the leaves being pinnately nerved (not trinerved), the number of lateral nerves 7 *–* 9 on each side, with domatia comprised of two tufts of white hairs (not 4 *–* 5(*–* 6) nerves and domatia absent), tertiary nerves strongly scalariform, leaf apex rounded or very shortly and broadly acuminate (versus reticulate, long-acuminate).

## DISTRIBUTION

Cameroon, S.W. Region, Rumpi Hills.

## HABITAT

Submontane forest with *Santiria trimera* (Burseraceae), *Garcinia smeathmannii* (Clusiaceae or Guttiferae) and *Carapa grandifolia* (now *C. oreophila* Kenfack, Meliaceae) (*D*.*W. Thomas* 10426); c. 1300 m alt.

## ETYMOLOGY

Named as a noun in apposition for the Rumpi Hills in S.W. Region, Cameroon, to which this species is restricted on current evidence.

## SPECIMENS EXAMINED. CAMEROON

S.W. Region, Rumpi Hills, near Madie River 4° 58’N, 9° 15’E, fr. 22 Feb 1995, *D*.*W*.*Thomas* 10426 (K holo. barcode K000875935; iso. EA, MA, US, YA); ibid fl.fr. 22 Feb 1995, *D*.*W*.*Thomas* 10442 (K barcode K001383574, YA).

## VERNACULAR NAMES & USES

None are recorded.

## CONSERVATION STATUS

Known from a single location, the higher altitudes of the Rumpi Hills, where it grows with the point endemic *Ocotea ikonyokpe* van der Werff. (Lauraceae, van der Werff 1996). The site of the type specimen collection of the last, *D*.*W. Thomas* 10456 is given as “l.5 km W of Madie River ford”. This species, in the identical habitat, with similar restricted range has been assessed for its global extinction risk status as threatened but is not yet published on iucnredlist.org (July 2022). *Microcos rumpi* is here assessed as Critically Endangered since only a single site is known, area of occupation is calculated as 4 km^2^ using the preferred IUCN grid cell size and extent of occurrence as the same. Plotting the grid reference of the two specimens on Google Earth shows that the site is outside the existing Rumpi Hills protected area, and examining historic imagery shows that between July 2008 and Jan. 2011, there is an increase in cleared areas of forest along the road c. 500m to the west of the grid reference, and also clearing distant from the road. This justifies an assessment of CR B1ab(iii) + B2ab(iii). This distinctive, huge tree has not been found in surveys elsewhere in the Cameroon Highlands and adjacent areas (Cheek 1992; Cheek *et al*. 1996; Cable & Cheek 1998; Cheek *et al*. 2000; Maisels *et al*. 2000; Chapman & Chapman 2001; Cheek *et al*. 2004; Harvey *et al*. 2004; Cheek *et al*. 2006; Cheek *et al*. 2010; Harvey *et al*. 2010; Cheek *et al*. 2011). Therefore, it may indeed be endemic to the Rumpi Hills.

## NOTES

The only known collections of *Microcos rumpi* appear to be connected with a transect made by Thomas (1995:37 *–* 39, table 5) in the Rumpi Hills. In that transect, 607 stems above 10 cm dbh (trunk diameter at 1.3 m above ground level) were recorded in five size classes and allocated to 80 different species (Thomas 1995: 37 *–* 39, table 5). The transect area was 1 Ha (1 km x 10 m) at elevation 1200 *–* 1350 m alt., starting at the Madie River crossing between Dikome and Madie, just to the East of the RHFR. Hilltops were reported to be dominated by *Santiria trimera* (Burseraceae) a well-known indicator species of submontane forest in Cameroon. *Microcos rumpi* (as “*Grewia coriacea”*) occurred 10 times in total, 5 stems (c. 1% of stems) in the 10 *–* 29 cm size classes, 1 stem in the 30 *–* 49 cm class (c. 1.5%), and 4 stems (c. 15%) in the 50 *–* 69 cm size class (Thomas 1995). Therefore, within the transect area, the species is not uncommon, since about 1 in every 60 stems above 10 cm dbh is *Microcos rumpi*. While the specimen location metadata relates to that given for the transect, the size class data given for the specimen (up to 80 cm dbh) exceeds that for any recorded in the transect, suggesting the specimen(s) were taken near to but outside the transect.

Since only two stems among the 607 of the Rumpi Hills transect are placed in the 90+ cm size class (Thomas 1995), at 80 cm dbh (*Thomas* 10426), *Microcos rumpi* must be among the most massive trees in its Rumpi Hills cloud forest habitat consistent with the height given of 35 *–* 40 m (*Thomas* 10426). Taking the average wood density of *Microcos* of 0.5 kg/dm^3^ from Dryad (Zanne *et al*. 2009) in the absence of African data derived from Asian species of *Microcos*, together with the metadata cited above from *Thomas* 10426, the equation for estimating the biomass of trees from moist tropical forests (Chave *et al*. 2005) gave an above ground biomass of such a tree as approximately 6500 kg, plus 1400 kg below ground (equation from Sierra *et al*. 2001), that is 7.9 metric tonnes per tree. That such an immense organism could remain unknown to science until now is remarkable if not without precedent.

At the time of specimen collection it was evident from the specimen metadata that the collector queried whether these specimens represented *Grewia coriacea* (*Microcos coriacea*) since the trees were so much larger than those of the species encountered at lower altitudes. But by the time the report was written (Thomas 1995) the specimens are unambiguously attributed to *Grewia coriacea* (*Microcos coriacea*).

While *Microcos rumpi* is most likely to share a most recent common ancestor with one of the two other Cameroonian tree *Microcos* species, *M. coriacea* and *M. magnifica* (see above), it shares a remarkable character with one of the most common and widespread Congolese species, *M. louisii*. Both species have leaf domatia comprised of two separate clusters of white hairs which interlock at the apex (Fig. 1C). The two species share other similarities in leaf shape but differ in the features indicated in Table 1. In addition, *M. louisii* is mainly confined to seasonally flooded and swamp forests of the Congo basin and is not recorded from cloud (submontane) forest in Cameroon.

**Table 1.** Major diagnostic characters separating *Microcos coriacea, M. louisii M. rumpi* and *M. magnifica*. Data from specimens at K and for the last species from Cheek (2017).

|  | *Microcos coriacea* | *Microcos louisii* | *Microcos rumpi* | *Microcos magnifica* |
| --- | --- | --- | --- | --- |
| Height (m) | 10 – 20 | 7 – 8 | 35 – 40 | 20 – 35 |
| Number of lateral nerves on each side of midrib | 4 – 5( 6) | 5 – 6 | (7 – )8 – 9( – 10) | 11 – 13 |
| Leaf-base | Acute to rounded | Truncate to rounded | (Rounded-) truncate(-cordate) | Truncate to abruptly cordate |
| Domatia | No | Yes | Yes | No |
| Nervation | Basally 3-nerved | Pinnately nerved | Pinnately nerved | Basally 3-nerved |
| Tertiary Nerves | Reticulate | Reticulate | Scalariform | Scalariform |
| Alt. range (m) | 0 – 500( – 1000) | 400 – 600 | c. 1300 | 750 – 1000 |
| Geography | Nigeria-Gabon | D.R.Congo | Rumpi Hills, Cameroon | Mt Kupe and Ebo Forest, Cameroon |

### The Rumpi Hills

The Rumpi Hills Forest Reserve (RHFR) is situated 80 km N of Mt Cameroon, 50 km W of Bakossi Mts and 15 km SE of Korup National Park. The highest point is given as Mt Rata, at 1800 m (https://en.wikipedia.org/wiki/Rumpi_Hills). Data on the Rumpi Hills is scarce. The most authoritative appears to be a study on land-use change by Beckline *et al*. (2018). They state that the Rumpi Hills exhibits two seasons; the dry season from November to April and the rainy season from May to October with annual rainfall that ranges between 4027–6368 mm. Mean monthly maximum temperatures in the dry season is estimated at 31.8°C and 18.2°C during the rainy season. The relative humidity is high during most of the year with minimum monthly values ranging between 78% and 90%. Ethnic groups are the Ngolo, Bima, and Balue. The Rumpi Hills Forest Reserve is given as 455 km^2^ and is kidney-shaped in outline. Google Earth imagery shows the forest canopy of RHFR to be largely intact, apart from oil palm plantations inside the far western boundary around the settlement of Ekumbako (4° 55’ 41.47”N, 80 56’ 18.43”E). However, the boundary shown on Google Earth may be inaccurate and this settlement may fall outside the boundary. Despite the name, most of RHFR apart from inside the northern and eastern boundary, is not aligned with the upland area of the Rumpi Hills, which lie to the east. RHFR is below 600 m and mostly below 300 *–* 400 m.

The highest settlement of the Rumpi Hills is Dikome Balue, the capital of the Rumpi Hills, which lies outside the RHFR. It is only 7 km S of the recorded location of *Microcos rumpi*. Smaller settlements, mostly above 1000 m alt. are scattered regularly through the hills proper, that occupy the area east and southeast of the RHFR. A recent study of changes in the Rumpi Hills between 2000 *–* 2014 based on satellite images and, importantly, on-the-ground surveys in 14 settlements around RHFR and 200 land-use change plots, reported that during the 14-year period, dense forest dropped to 90.2% while settlements increased from 744.6 to 2148.8 hectares in 2014. Also, farmlands increased by 18.25% representing a change from 9,400 to 11,117 Ha (Beckline *et al*. 2018). Despite this, observation of Google Earth imagery dating from 2015 (accessed 17 July 2022) shows large areas of forest with intact canopy. The presence of the range-restricted endemics *Ocotea ikonyokpe* (see above), and, also outside the boundary of the RHFR, to the north, the point endemic *Ledermanniella prasina* J. J. Schenk & D. W. Thomas (Podostemaceae, Schenck & Thomas 2004), now joined by a third Rumpi Hills endemic species, *Microcos rumpi* (this paper) are indicators that the Rumpi Hills are a centre of plant diversity. That the numbers of recorded endemic plant species is so low compared with the neighbouring Bakossi Mts and Mt Kupe (Cheek *et al*. 2004) is undoubtedly down to the comparatively very low levels of botanical survey and identification in the Rumpi Hills. Undoubtedly many additional new species to science, including Rumpi Hill endemic species, will be brought to light if botanical collections and identifications are resumed in the Rumpi Hills while natural habitat remains.

## Discussion

*Microcos rumpi* is the latest in a long line of discoveries of new species to science from the cloud forest habitats of the Cameroon Highlands. These species vary from epiphytic herbs e.g. *Impatiens frithi* Cheek (Balsaminaceae, Cheek & Csiba 2002) and *I. etindensis* Cheek & Eb. Fisch. (Balsaminaceae, Cheek & Fischer 1999) to terrestrial herbs e.g. *Brachystephanus kupeensis* I. Darbysh. & Champl. (Acanthaceae, Champluvier & Darbyshire 2009) and *Isoglossa dispersa* I. Darbysh. (Darbyshire *et al*. 2011), rheophytes e.g. *Ledermaniella onanae* Cheek (Cheek 2003) and *Saxicolella ijim* Cheek (Podostemaceae; Cheek *et al*. 2022a) to achlorophyllous mycotrrophs e.g. *Kupea martinetugei* Cheek & S.A.Williams (Triuridaceae, Cheek *et al*. 2003) and *Afrothismia amietii* Cheek (Cheek 2007, Thismiaceae) and understorey shrubs and small trees e.g. *Kupeantha kupensis* Cheek (Rubiaceae, Cheek *et al*. 2018a) and *Psychotria darwiniana* Cheek (Rubiaceae, Cheek *et al*. 2009), to canopy trees, e.g. *Vepris zapfackii* Cheek & Onana, *Deinbollia onanae* Cheek, and *Vepris onanae* Cheek (Cheek & Onana 2021; Cheek *et al*. 2021a; 2022b)

Most of these new species were first described as point or near-endemics, but several have, with more research, been found to be more widespread e.g. *Tricalysia elmar* Cheek (Rubiaceae, Cheek *et al*. 2020a) now known to extend from Mt Kupe and Bali-Ngemba to e.g. the Rumpi Hills (Lachenaud *et al*. 2013), *Coffea montekupensis* Stoff. (Stoffelen *et al*. 1997) first thought to be endemic to Mt Kupe is now known to occur in the Tofala Sanctuary (Lebialem Highlands, Harvey *et al*. 2010) and *Oxyanthus okuensis* Cheek & Sonké (Rubiaceae, Cheek & Sonké 2000) initially thought to be endemic to Mt Oku is now known to extend to Tchabal Mbabo (Lachenaud *et al*. 2013). It is to be hoped that the range of *Microcos rumpi* will be similarly extended by future research and its extinction risk assessment consequently reduced.

## Conclusion

It is important to uncover the existence of previously unknown plant species as soon as possible and to formally name them. Until this is done, they are invisible to science and the possibility of their being assessed for their conservation status and appearing on the IUCN Red List is greatly reduced (Cheek *et al*. 2020b), limiting the likelihood that they will be proposed for conservation measures, and that such measures will be accepted. Although there are exceptions (Cheek & Etuge 2009; Cheek *et al*. 2019a), most new plant species to science are highly range-restricted, making them almost automatically threatened (Cheek *et al*. 2020b).

Only 7.2% of the 369,000 flowering plant species (the number is disputed) known to science have been assessed on the IUCN Red List. (Bachman *et al*. 2019; Nic Lughadha *et al*. 2016; 2017). However, the vast majority of plant species still lack assessments on the Red List (Nic Lughadha *et al*. 2020). Fortunately, Cameroon has a plant Red Data book (Onana & Cheek 2011), which details 815 threatened species, but it needs updating. Thanks to the Global Tree Assessment (BGCI 2021) many of the world’s tree species have now been assessed. The State of the World’s Trees concluded that the highest proportion of threatened tree species is found in Tropical Africa, and that Cameroon has the highest number (414) of threatened tree species of all tropical African countries (BGCI 2021). This will be further increased by the addition of *Microcos rumpi*.

Concerns about global plant species extinctions are increasing as the biodiversity crisis continues. In Cameroon, the lowland forest species are *Oxygyne triandra* Schltr., *Afrothismia pachyantha* Schltr. have been considered extinct for some years (Cheek & Williams 1999, Cheek *et al*. 2018b, Cheek *et al*. 2019b). Similarly, *Pseudohydrosme bogneri* Cheek & Moxon-Holt and *P. buettneri* Engl. are now considered extinct in lowland forest in neighbouring Gabon (Moxon-Holt & Cheek 2020; Cheek *et al*. 2021b). However, submontane (cloud) forest species are now also being recorded as extinct in Cameroon, such as the well-documented case of *Vepris bali* Cheek *et al*. 2018c), and the only recently discovered *Monanthotaxis bali* Cheek (Annonaceae, Cheek *et al*. 2022c). Global species extinctions are being recorded from across Africa, from West (e.g. *Inversodicraea pygmaea* G. Taylor and *Saxicolella deniseae* Cheek in Guinea (Cheek *et al*. 2017; 2022a)) to East (e.g. *Kihansia lovettii* Cheek and *Vepris* sp A of FTEA in Tanzania (Cheek 2004; Cheek & Luke 2022)).

If such extinctions are to cease, or more realistically, to be slowed, improved conservation prioritisation programmes are needed to firstly determine the most important plant areas for conservation (Darbyshire *et al*. 2017) and secondly to implement protection with local communities and authorities through well-drawn up management plans and resourcing. Cultivation and seedbanking of species at risk of extinction, if feasible, are important fall-back strategies, but conservation of species in their natural habitat must be the first priority.

The survival of *Microcos rumpi* may not seem urgent at present because while data on habitat loss around the Rumpi Hills shows a steady increase (Beckline *et al*. 2017), there are also still large tracts of habitat seemingly intact where the species may survive (Google Earth imagery, accessed July 2022). However, scenarios can change very rapidly, as with *Saxicolella deniseae* which when collected for the first time in 2018 was not considered to be at risk of extinction, but which appears to have become globally extinct in 2020 or 2021, before it was published (Cheek *et al*. 2022a).

## Acknowledgements

This paper was completed as part of the Cameroon TIPAs (Tropical Important Plant Areas) project at RBG, Kew, which is supported by Players of Peoples Postcode Lottery. We thank Lydia Burns and Penny Appelbe of Kew’s Foundation for making this possible. This paper is a result of the partnership between RBG, Kew and IRAD-National Herbarium of Cameroon, Yaoundé and we thank the late Dr Benoît Satabié, Drs Gaston Achoundong, Florence Ngo Ngwe, Eric Nana, Jean Lagarde Betti, the current and former directors or acting Directors, of IRAD-National Herbarium of Cameroon, Yaoundé, for expediting the collaboration between our two institutes. Dr Duncan Thomas provided helpful comments to the Rumpi Hills TIPA datasheet compiled by Dr Bruce Murphy, giving useful context to the discussion of this paper. Rosemary Lomer, volunteer with the Cameroon TIPAs project, uncovered the specimens of what turned out to be the new species, and brought them to attention: without her this species would still remain unknown to science. Janis Shillito typed the manuscript. Two anonymous reviewers are thanked for constructively reviewing an earlier version of this paper.

The authors declare that they have no conflict of interest.

## References

African Plant Database (version 3.4.0). (continuously updated). Conservatoire et Jardin botaniques de la Ville de Genève and South African National Biodiversity Institute, Pretoria, “Retrieved [March 2017]”, from <http://www.ville-ge.ch/musinfo/bd/cjb/africa/>.

Bachman, S.P., Field, R., Reader, T., Raimondo, D., Donaldson, J., Schatz, G.E. and Lughadha, E.N. (2019). Progress, challenges and opportunities for Red Listing. Biological Conservation 234: 45–55. https://doi.org/10.1016/j.biocon.2019.03.002

Bayer, C., Fay, M.F., De Bruijn, A.Y., Savolainen, V., Morton, C.M., Kubitzki, K., Alverson, W.S., Chase, M.W. (1999). Support for an expanded family concept of Malvaceae within a recircumscribed order Malvales: a combined analysis of plastid atpB and rbcL DNA sequences. Botanical Journal of the Linnean Society 129: 267–303. https://doi.org/10.1111/j.1095-8339.1999.tb00505.x

Bayer, C., Kubitzki, K. (2003). Malvaceae. In: Kubitzki, K., Bayer, C. (eds), The families and genera of vascular plants 5: flowering plants dicotyledons: 225–311. Springer-Verlag, Berlin.

Beckline, M., Yujun, S., Etongo, D., Saeed, S., & Mannan, A. (2018). Assessing the drivers of land use change in the Rumpi hills forest protected area, Cameroon. Journal of Sustainable Forestry 37(6), 592–618. https://doi.org/10.1080/10549811.2018.1449121

Beentje, H. & Cheek, M. (2003). Glossary. In Beentje, (ed.), Flora of Tropical East Africa. Balkema, Lisse, Netherlands.

BGCI (2021). State of the World’s Trees. BGCI, Richmond, UK.

Brunken, U. & Muellner, A.N. (2012). A New Tribal Classification of Grewioideae (Malvaceae) Based on Morphological and Molecular Phylogenetic Evidence. Systematic Botany 37(3):699–711. http://www.bioone.org/doi/full/10.1600/036364412X648670

Burret, M. (1926). Beiträge zur Kenntnis der Tiliaceae I. Notizblatt des Königlich-Botanischen Gartens und Museums zu Berlin-Dahlem 9: 592–880.

Cable, S. & Cheek, M. (1998). The Plants of Mt Cameroon, a Conservation Checklist. Royal Botanic Gardens, Kew.

Champluvier, D. & Darbyshire, I. (2009). A revision of the genera Brachystephanus and Oreacanthus (Acanthaceae) in tropical Africa. Systematics and Geography of Plants 79(2):115–192. https://doi.org/10.2307/25746

Chapman, J. & Chapman, H. (2001). The Forests of Taraba and Adamawa States, Nigeria an Ecological Account and Plant Species Checklist. University of Canterbury: Christchurch, New Zealand. pp. 221.

Chattaway, M.M. (1934). Anatomical evidence that Grewia and Microcos are distinct genera. Tropical Woods 38: 9–11

Chave, J., Andalo, C., Brown, S., Cairns, M.A., Chambers, J.Q., Eamus, D., Folster, H., Fromard, F., Higuchi, N., Kira, T., Lescure, J.P., Nelson, B.W., Ogawa, H., Puig, H., Riéra, B., Yamakura, T. (2005). Tree allometry and improved estimation of carbon stocks and balance in tropical forests. Oecologia 145: 87–99. https://doi.org/10.1007/s00442-005-0100-x

Cheek, M. (1992). A Botanical Inventory of the Mabeta-Moliwe Forest. Royal Botanic Gardens, Kew.

Cheek, M. (2003). A new species of Ledermanniella (Podostemaceae) from western Cameroon. Kew Bulletin 58: 733–737. https://doi.org/10.2307/4111153

Cheek, M. (2004). Kupeaeae, a new tribe of Triuridaceae from Africa. Kew Bulletin 58: 939–949. https://doi.org/10.2307/4111207

Cheek, M. (2007). Afrothismia amietii (Burmanniaceae), a new species from Cameroon. Kew Bull. 61: 605–607. http://www.jstor.org/stable/20443306 https://doi.org/10.7717/peerj.4137

Cheek, M. (2017). Microcos magnifica (Sparrmanniaceae) a new species of cloudforest tree from Cameroon. PeerJ 5:e4137

Cheek, M. & Cable, S. (1997). Plant Inventory for conservation management: the Kew-Earthwatch programme in Western Cameroon, 1993–96, pp. 29–38 in Doolan, S. (Ed.) African Rainforests and the Conservation of Biodiversity, Earthwatch Europe, Oxford.

Cheek, M. & Csiba, L. (2002). A new epiphytic species of Impatiens (Balsaminaceae) from western Cameroon. Kew Bull. 57: 669–674. https://doi.org/10.2307/4110997

Cheek, M. & Etuge, M. (2009). A new submontane species of Deinbollia (Sapindaceae) from Western Cameroon and adjoining Nigeria. Kew Bull. 64: 503–508. https://doi.org/10.1007/s12225-009-9132-4

Cheek, M. & Fischer, E. (1999). A tuberous and epiphytic new species of Impatiens (Balsaminaceae) from Southwest Cameroon. Kew Bull. 54: 471–475. https://doi.org/10.2307/4115828

Cheek, M. & Luke, W.R.Q. (2022). A taxonomic synopsis of unifoliolate continental African Vepris (Rutaceae) with three new threatened forest tree species from Kenya and Tanzania and two possibly extinct. bioRix. doi: https://doi.org/10.1101/2022.07.16.500287

Cheek, M. & Onana, J.M. (2021). The endemic plant species of Mt Kupe, Cameroon with a new Critically Endangered cloud-forest tree species, Vepris zapfackii (Rutaceae). Kew Bull 76, 721–734 https://doi.org/10.1007/s12225-021-09984-x

Cheek, M. & Sonké, B. (2000). A new species of Oxyanthus (Rubiaceae-Gardeniinae) from western Cameroon. Kew Bulletin 55: 889–893. https://doi.org/10.2307/4113634

Cheek, M. & Williams, S. (1999). A Review of African Saprophytic Flowering Plants. In: Timberlake, Kativu eds. African Plants. Biodiversity, Taxonomy & Uses. Proceedings of the 15th AETFAT Congress at Harare. Zimbabwe, 39–49.

Cheek, M., Achoundong, G., Onana, J-M., Pollard, B., Gosline, G., Moat, J., Harvey, Y.B. (2006). Conservation of the Plant Diversity of Western Cameroon. In: Ghazanfar SA, H.J. Beentje (eds). Proceedings of the 17th AETFAT Congress, Addis Ababa. Ethiopia, 779–791.

Cheek, M., Alvarez-Agiurre, M.G., Grall, A., Sonké, B., Howes, M-J.R., Larridon, I. (2018a). Kupeantha (Coffeeae, Rubiaceae), a new genus from Cameroon and Equatorial Guinea. PLoS ONE 13: 20199324. https://doi.org/10.1371/journal.pone.0199324

Cheek, M., Cable, S., Hepper, F.N., Ndam, N., Watts, J. (1996). Mapping plant biodiversity on Mt. Cameroon. In: Maesen, Burgt Rooy eds. The Biodiversity of African Plants (Proceedings XIV AETFAT Congress. Cameroon: Kluwer, 110–120. https://doi.org/10.1007/978-94-009-0285-5_16

Cheek, M., Causon, I., Tchiengue, B. & House, E. (2020a). Notes on the endemic cloud forest plants of the Cameroon Highlands and the new, Endangered, Tricalysia elmar (Coffeeae-Rubiaceae). Plant Ecology and Evolution 153: 167–176 https://doi.org/10.5091/plecevo.2020.1661

Cheek, M., Corcoran, M. & Horwath, A. (2009b). Four new submontane species of Psychotria (Rubiaceae) with bacterial nodules from western Cameroon. Kew Bull. 63: 405–418. https://doi.org/10.1007/s12225-008-9056-4

Cheek, M., Darbyshire, I. & Onana, J.-M. (2022c). Monanthotaxis bali (Annonaceae) a new Critically Endangered (possibly extinct) montane forest treelet from Bali Ngemba, Cameroon. BioRxiv https://doi.org/10.1101/2022.07.04.498636

Cheek, M., Etuge, M. & Williams, S. (2019b). Afrothismia kupensis sp. nov. (Thismiaceae), Critically Endangered, with observations on its pollination and notes on the endemics of Mt Kupe, Cameroon. Blumea 64: 158–164. https://doi.org/10.3767/blumea.2019.64.02.06

Cheek M, Feika A, Lebbie A, Goyder D, Tchiengue B, Sene O, P. Tchouto P, Burgt X. (2017). A synoptic revision of Inversodicraea (Podostemaceae). Blumea 62, 2017: 125–156. https://doi.org/10.3767/blumea.2017.62.02.07

Cheek, M., Gosline, G. & Onana, J.M. (2018c). Vepris bali (Rutaceae), a new critically endangered (possibly extinct) cloud forest tree species from Bali Ngemba, Cameroon. Willdenowia 48: 285–292. https://doi.org/10.3372/wi.48.48207

Cheek, M., Harvey, Y.B., Onana, J-M. (2010). The Plants of Dom. Bamenda Highlands, Cameroon: A Conservation Checklist. Royal Botanic Gardens, Kew.

Cheek M, Harvey Y, Onana J-M. (2011). The Plants of Mefou Proposed National Park. Yaoundé, Cameroon: A Conservation Checklist. Royal Botanic Gardens, Kew.

Cheek, M., Hatt, S., & Onana, J. M. (2022b). Vepris onanae (Rutaceae), a new Critically Endangered cloud-forest tree species, and the endemic plant species of Bali Ngemba Forest Reserve, Bamenda Highlands Cameroon. Kew Bull., 1–15. https://doi.org/10.1007/s12225-022-10020-9

Cheek, M., Molmou, D., Magassouba, S., & Ghogue, J. P. (2022a). Taxonomic revision of Saxicolella (Podostemaceae), African waterfall plants highly threatened by Hydro-Electric projects. Kew Bull., 1–31. https://doi.org/10.1007/s12225-022-10019-2

Cheek, M., Nic Lughadha, E., Kirk, P., Lindon, H., Carretero, J., Looney, B., Douglas, B., Haelewaters, D., Gaya, E., Llewellyn, T., Ainsworth, M., Gafforov, Y., Hyde, K., Crous, P., Hughes, M., Walker, B.E., Forzza, R.C., Wong, K.M., Niskanen, T. (2020b). New scientific discoveries: plants and fungi. Plants, People Planet 2: 371–388. https://doi.org/10.1002/ppp3.10148 https://doi.org/10.7717/peerj.11036

Cheek, M., Onana, J.M., Chapman, H.M. (2021a). The montane trees of the Cameroon Highlands, West-Central Africa, with Deinbollia onanae sp. nov. (Sapindaceae), a new primate-dispersed, Endangered species. PeerJ 9:e11036

Cheek, M., Onana, J-M., Pollard, B.J. (2000). The Plants of Mount Oku and the Ijim Ridge, Cameroon, a Conservation Checklist. Royal Botanic Gardens, Kew.

Cheek, M., Onana, J-M., Yasuda, S., Lawrence, P., Ameka, G., Buinovskaja G. (2019a). Addressing the Vepris verdoorniana complex (Rutaceae) in West Africa, with two new species. Kew Bull. 74: 53. https://doi.org/10.1007/S12225-019-9837-Y

Cheek, M., Pollard, B.J., Darbyshire, I, Onana, J.M. & Wild, C. (2004). The Plants of Kupe, Mwanenguba and the Bakossi Mts, Cameroon. A Conservation Checklist. Royal Botanic Gardens, Kew.

Cheek, M., Tchiengué, B., van der Burgt, X. (2021b). Taxonomic revision of the threatened African genus Pseudohydrosme Engl. (Araceae), with P. ebo, a new, critically endangered species from Ebo, Cameroon. PeerJ 9:e10689 https://doi.org/10.7717/peerj.10689.

Cheek, M., Tsukaya, H., Rudall, P.J., Suetsugu, K. (2018b). Taxonomic monograph of Oxygyne (Thismiaceae), rare achlorophyllous mycoheterotrophs with strongly disjunct distribution. PeerJ 6: e4828. https://doi.org/10.7717/peerj.4828

Cheek, M., Williams, S. & Etuge, M. (2003). Kupea martinetugei, a new genus and species of Triuridaceae from western Cameroon. Kew Bulletin 58: 225–228. https://doi.org/10.2307/4119366

Chung, R.C.K. (2002). Leaf epidermal micromorphology of Grewia L. and Microcos L. (Tiliaceae) in Peninsular Malaysia and Borneo. Gardens’ Bulletin Singapore 54: 263–286.

Chung, R.C.K. (2003). New taxa and new combinations of Microcos (Tiliaceae) from Peninsular Malaysia and Borneo. Kew Bulletin 58: 329–349. https://doi.org/10.2307/4120619

Chung, R.C.K. (2006). A revision of Grewia (Malvaceae-Grewioideae) in Peninsular Malaysia and Borneo. Edinburgh Journal of Botany 62: 1–27. https://doi.org/10.1017/s0960428606000011

Chung, R.C.K., Lim, S.C., Lim, A.L., Soepadmo, E. (2005b). Wood anatomy of Grewia and Microcos from Peninsular Malaysia and Borneo. Journal of Tropical Forest Science 17: 175–196.

Chung, R.C.K., Soepadmo, E., Lim, A.L. (2003). The significance of pollen morphology in the taxonomy of Grewia and Microcos (Tiliaceae) in Peninsular Malaysia and Borneo. Gardens’ Bulletin Singapore 55: 239–256.

Chung, R.C.K., Soepadmo E, Lim, A.L. (2005a). A synopsis of the Bornean species of Microcos L. (Tiliaceae). Gardens’ Bulletin Singapore 57: 103–132.

Chung, R.C.K. & Soepadmo, E. (2011). Taxonomic revision of the genus Microcos (Malvaceae-Grewioideae) in Peninsular Malaysia and Singapore. Blumea 56: 273–299. http://dx.doi.org/10.3767/000651911X619704

Darbyshire, I., Anderson, S., Asatryan, A., Byfield, A., Cheek, M., Clubbe, C., Ghrabi, Z., Harris, T., Heatubun, C. D., Kalema, J., Magassouba, S., McCarthy, B., Milliken, W., Montmollin, B. de, Nic Lughadha, E., Onana, J.M., Saidou, D., Sarbu, A., Shrestha, K. & Radford, E. A. (2017). Important Plant Areas: revised selection criteria for a global approach to plant conservation. Biodivers. Conserv. 26: 1767–1800. https://doi.org/10.1007/s10531-017-1336-6.

Darbyshire, I., Pearce, L. & Banks, H. (2011). The genus Isoglossa (Acanthaceae) in west Africa. Kew Bulletin 66 (3): 425–439. http://dx.doi.org/10.1007/s12225-011-9292-x

Govaerts, RHA, Belyaeva I, Hartley H & Lindon H. (continuously updated). Microcos Burm. ex L.in: Plants of the World Online. Kew Science. powo.science.kew.org/taxon/urn:lsid:ipni.org:names:328157–2

Harvey, Y.B., Pollard, B.J., Darbyshire, I., Onana, J.-M., Cheek, M. (2004). The Plants of Bali Ngemba Forest Reserve. Cameroon: A Conservation Checklist. Royal Botanic Gardens, Kew.

Harvey, Y.B., Tchiengue, B., Cheek, M. (2010). The Plants of the Lebialem Highlands, a Conservation Checklist. Royal Botanic Gardens, Kew.

Heywood, V.H., Brummitt, R. K., Culham, A. & Seburg, O. (eds.) (2007). Flowering Plant Families of the World. Royal Botanic Gardens, Kew.

IPNI (continuously updated). The International Plant Names Index. http://ipni.org/.

IUCN. 2012. IUCN Red List Categories and Criteria: Version 3.1. Second edition. Gland, Switzerland and Cambridge, UK: IUCN. Available from: http://www.iucnredlist.org/ (accessed: 01/2017).

Lachenaud, O., Droissart, V., Dessein, S., Stévart, T., Simo, M., Lemaire, B., Taedoumg, H. and Sonké, B. (2013). New records for the flora of Cameroon, including a new species of Psychotria (Rubiaceae) and range extensions for some rare species. Plant Ecology and Evolution 146: 121–133. https://doi.org/10.5091/plecevo.2013.632

Linnaeus C. 1753. Species plantarum. Salvii, Stockholm.

Linnaeus C. 1767. Systema naturae ed. 12, 1, 2. Salvii, Stockholm.

Maisels, F.M., Cheek, M., Wild, C. (2000). Rare plants on Mt Oku summit, Cameroon. Oryx 34: 136–140. https://doi.org/10.1017/s0030605300031057.

Masters, M. T. (1868). Tiliaceae. In Oliver, D. (ed), Flora of Tropical Africa 1: 240–268. L. Reeve & Co., London

Moxon-Holt, L. & Cheek, M. (2020). Pseudohydrosme bogneri sp. nov. (Araceae), a spectacular Critically Endangered (Possibly Extinct) species from Gabon, long confused with Anchomanes nigritianus. BioRxiv (pre-print) https://doi.org/10.1101/2021.03.25.437040

Nic Lughadha, E., Bachman, S.P., Govaerts, R. (2017). Plant fates and states: Response to Pimm & Raven. Trends in Ecology and Evolution 32: 887–889

Nic Lughadha, E., Bachman, S.P., Leão, T.C., Forest, F., Halley, J.M., Moat, J., Acedo, C., Bacon, K.L., Brewer, R.F., Gâteblé, G., Gonçalves, S.C., Govaerts, R., Hollingsworth, P.M., Krisai-Greilhuber, I., de Lirio, E.J., Moore, P.G.P., Negrão, R., Onana, J.M., Rajaovelona, L.R., Razanajatovo, H., Reich, P.B., Richards, S.L., Rivers, M.C., Amanda Cooper, A., Iganci, J., Lewis, G.L., Smidt, E.C., Antonelli, A., Mueller, G.M. & Walker, B.E. (2020). Extinction risk and threats to plants and fungi. Plants, People, Planet 2: 389–408. https://doi.org/10.1098/rstb.2017.0402

Nic Lughadha, E., Govaerts, R., Belyaeva, I., Black, N., Lindon, H. Allkin, R. Magill R.E. & Nicolson. (2016). Counting counts: Revised estimates of numbers of accepted species of flowering plants, seed plants, vascular plants and land plants with a review of other recent estimates. Phytotaxa 272: 82–88. https://doi.org/10.11646/phytotaxa.272.1.5

Onana, J.-M. & Cheek, M. (2011). The Red Data Book, Plants, of Cameroon. Royal Botanic Gardens, Kew

Schenk, J. J., & Thomas, D. W. (2004). A New Species of Ledermanniella (Podostemaceae) from Cameroon. Novon, 14(2), 227–232. http://www.jstor.org/stable/3393321

Sierra, C., del Valle, J. & Orrego, S. (2001). Biomasa de raíces en bosques primarios y secundarios del área de influencia de la Central Hidroeléctrica Porce II. Trabajo de grado ingeniero forestal. Medellín, Colombia. Universidad Nacional de Colombia. https://repositorio.unal.edu.co/handle/unal/11262

Stoffelen, P., Cheek, M., Bridson, D., Robbrecht, E. (1997.) A new species of Coffea (Rubiaceae) and notes on Mt Kupe (Cameroon). Kew Bulletin 52(4): 989–994. https://doi.org/10.2307/3668527

Thiers, B. (continuously updated). Index Herbariorum: A global directory of public herbaria and associated staff. New York Botanical Garden’s Virtual Herbarium. [continuously updated]. Available from: http://sweetgum.nybg.org/ih/ (accessed: July 2022).

Thomas, D.W. (1995). Botanical Survey of the Rumpi Hills and Nta Ali. With special focus on the submontane zone above 1000m elevation. A report to GTZ. Corvallis, Oregon.

Van der Werff, H. (1996). Ocotea ikonyokpe, a new species of Lauraceae from Cameroon. Novon 6:460–462. https://doi.org/10.2307/3392056

Whitehouse, C., Andrews, S., Verdcourt, B. & Cheek, M. (2001). Tiliaceae. Flora of Tropical East Africa. Royal Botanic Gardens, Kew.

Wilczek, R. (1963a). Tiliaceae in Boutique, R. (ed.) Flore du Congo Belge et du Ruanda-Urundi, Spermatophytes 10: 1–91

Wilczek, R. (1963b). Novitates africanae VIII: Tiliaceae. Bulletin du Jardin botanique de l’État a Bruxelles 33(4): 459–471. https://doi.org/10.2307/3667378,

Zanne, A.E., Lopez-Gonzalez, G., Coomes, D.A., Ilic, J., Jansen, S., Lewis, S.L., Miller, R.B., Swenson, N.G., Wiemann, M.C. and Chave, J. (2009). Global wood density database. Dryad. https://doi.org/10.5061/dryad.234.

